# Bows and swords: why bacteria carry short and long-range weapons

**DOI:** 10.1101/2022.10.13.512033

**Authors:** Sean C. Booth, William P.J. Smith, Kevin R. Foster

## Abstract

Bacteria possess a diverse range of mechanisms for inhibiting competitors, including bacteriocins, tailocins, the type VI secretion system, and contact-dependent inhibition. Why bacteria have evolved such a wide array of weapon systems remains a mystery. Here we develop an agent-based model to compare short-range weapons that require cell-cell contact, with long-range weapons that rely on diffusion. Our models predict that contact weapons are useful when an attacking strain is outnumbered, facilitating invasion and establishment. By contrast, ranged weapons tend to only be effective when attackers are abundant. We test our predictions with the opportunistic pathogen *Pseudomonas aeruginosa*, which naturally carries multiple weapons, including contact-dependent inhibition (CDI) and diffusing tailocins. As predicted, short-range CDI functions better at low frequency, while long-range tailocins require high frequency and cell density to function effectively. Head-to-head competitions between the two weapon types further support our predictions: a tailocin attacker only defeats CDI when it is numerically dominant, but then we find it can be devastating. Finally, we show that the two weapons work well together when one strain employs both. We conclude that short and long-range weapons serve different functions and allow bacteria to fight both as individuals and as a group.

## Introduction

One of the most striking illustrations of Darwin’s ‘struggle for existence’^1^ is the evolution of weaponry^2,3^. Weapons – traits that evolved, at least in part, to injure and harm competitors – have evolved many times in animals, with examples in groups as diverse as trilobites, insects, mammals and dinosaurs^2^. Bacteria are a second group of organisms that commonly evolve weapons^3–5^. Many clinical antibiotics were first isolated from bacteria that release them into the environment to inhibit competitors^6–10^. Ribosomally-synthesized bacteriocins are deployed in a similar manner and include both chemical toxins and phage-tail derived tailocins which physically punch holes in competitors^11^. Bacteria also deploy close-range weapons that require cell-cell contact. Examples include type VI secretion systems, which fire toxin-laden needles into competing cells^12^, and contact-dependent inhibition systems, which are toxin-loaded filaments anchored to the outside of the cell^13^. The diversity of weapons seen in bacteria, therefore, certainly rivals that seen in animals. However, there is a notable difference between the two groups. Animal species tend to carry a single weapon type – e.g. horns *or* antlers *or* tusks – with a few potential exceptions (the dinosaur *Ankylosaurus magniventris* had both horns and a bludgeoning tail club)^2^. By contrast, bacteria commonly carry multiple types of weapon^3^.

*Pseudomonas aeruginosa* is a problematic opportunistic pathogen, due to its ability to withstand numerous antibiotics^14^. Alongside its defensive capacity, this species is a striking illustration of how many weapons bacteria can carry. *P. aeruginosa* produces multiple bacteriocins and toxic small molecules, which serve as long range weapons^15^. In addition, it can deploy contact-dependent inhibition and up to three type VI secretion systems as short-range weapons^16^. More generally, among species whose weapons have been characterized in detail, many carry both short and long-range weapons, including strains of *Bacteroides fragilis*^17,18^, *Pectobacterium carotovorum*^19,20^, *Burkholderia cepacia*^21,22^, *Chromobacterium violaceum*^23,24^ and *Myxococcus xanthus*^25,26^. What is the evolutionary basis for the prevalence of multiple weapons? One argument is that bacteria are simply more aggressive than other species such as animals, and that this favours the simultaneous use of multiple weapons. Consistent with this hypothesis, experiments suggest that bacteria engage in combat much more regularly than animals^3^, which typically avoid using their weapons^2^. However, a general increase in aggression does not explain why bacteria carry multiple *types* of weapons, as opposed to simply just investing more in a single type. We hypothesized that bacteria carry multiple weapons because they serve different functions during competition. We further reasoned that this explanation is most compelling for weapons that function at different ranges, a factor which can strongly influence the outcome of bacterial contests^27^. We therefore sought to test this hypothesis by performing a direct comparison of the competitive benefits of short versus long-range weapons.

We first employ a realistic agent-based model of bacterial competition that has previously been used to understand the evolutionary function of single weapons^28,29^. The power of this framework is that it allows one to rapidly study a wide variety of competition scenarios with relative ease, while being realistic enough to generate focused predictions for empirical testing. The model predicts that short- and long-range weapons do indeed have the potential to serve different functions. We test these predictions by genome editing *P. aeruginosa* strain PAO1 to generate strains that are susceptible to its own short- and long-range weapons (contact-dependent inhibition and tailocins, respectively). This approach allows us to directly compare weapons’ functioning in a controlled genetic background. In support of the modelling, we find that short and long-range weapons can provide cells with different advantages during combat. Contact weapons can be effective when an attacking strain is outnumbered, facilitating invasion and establishment. By contrast, ranged weapons are only effective when attackers are abundant, but here they prove to be a devastating form of attack.

## Results

### An agent-based model of short and long-range weapons

We first employ an established computational model where each cell is simulated as an individual agent (hence, ‘agent’ or ‘individual-based’ model, Methods)^28–32^. Bacterial cells are seeded onto a two-dimensional surface where they grow and divide to fill a vertical space, as can occur in a biofilm for example, or at the intestinal mucosa of a mammalian host. Cells interact with each other both physically – pushing and displacing each other as they grow and collide – and chemically, by producing toxins that can inhibit strains of a different genotype (Figure 1, Movie S1). When cells reach a specified height they are removed, mimicking dispersal or shedding from the top of the community. Bacterial weapons can vary substantially in a wide range of properties, including method of delivery (short or long-range), quantity produced, deadliness, cost of production and, of course, the strain or species where they are found^3^. This can make like-for-like comparisons of different weapons empirically challenging. However, with a model one can precisely define, and systematically vary, such properties of bacterial weapons and study their effects (Methods). The model is also spatially explicit, which is particularly important for contact-based weapons whose action depends upon the occurrence of physical contacts between cells^28,29^.

**Figure 1:**
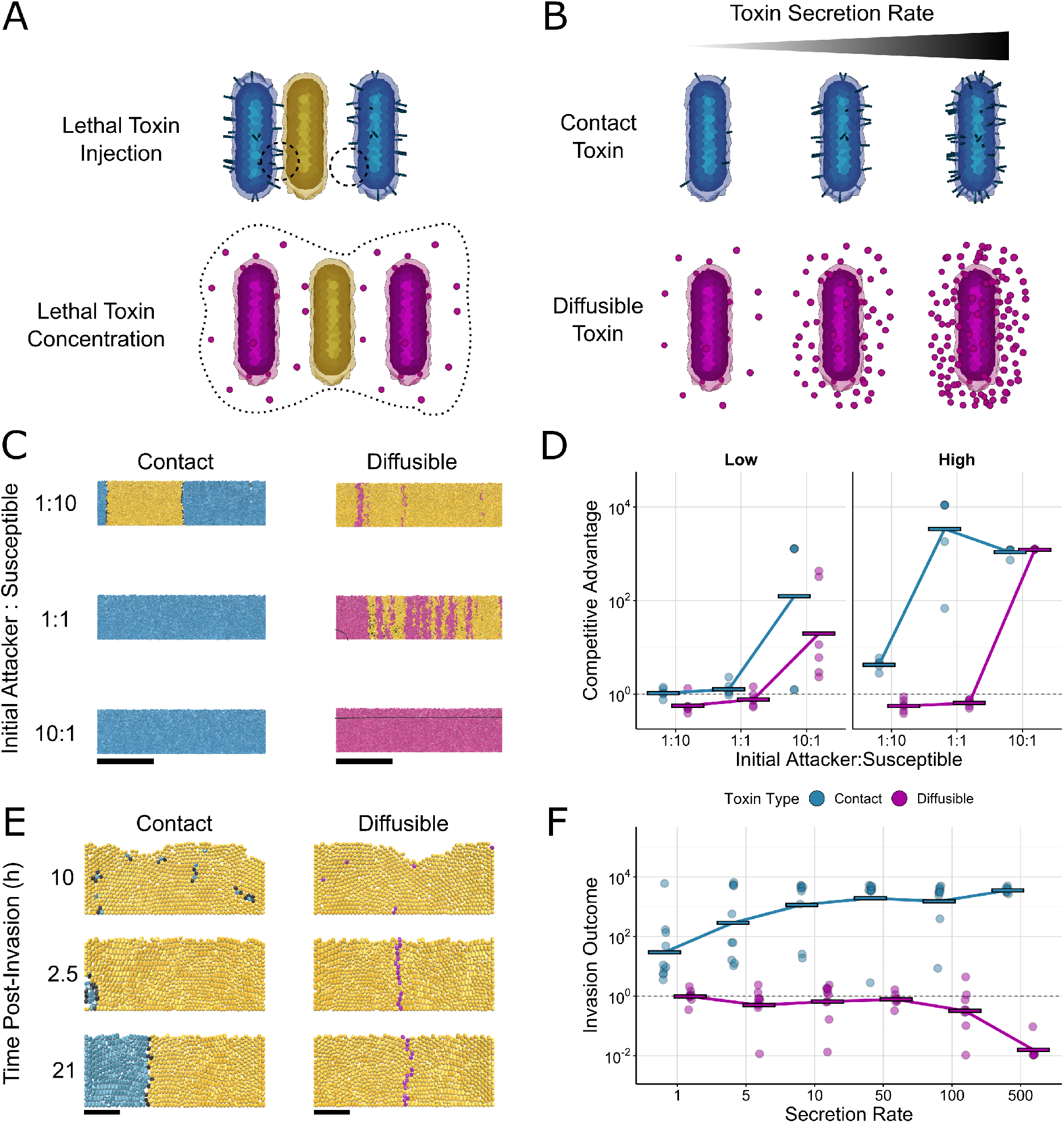
Agent-based modelling predicts that contact weapons are more robust to changes in frequency, density, and secretion rate. **A** Contact toxins (top): producing cells can deliver toxins to neighbouring cells. If a susceptible cell (yellow) is within range, the toxin is injected (left dashed circle) and the susceptible cell dies; otherwise the toxin is wasted (right dashed circle). Diffusing toxins (bottom): when the local concentration of a diffusible toxin exceeds a threshold (within dashed line), susceptible cells die. **B** Cells secrete toxins, incurring a growth-rate penalty proportional to the amount of toxin being secreted (secretion rate). **C** Snapshots of competition outcomes for attackers with contact-dependent toxins (blue cells, left column) or diffusible toxins (magenta cells, right column) Unarmed susceptibles (yellow cells) die upon lethal toxin exposure (black cells). The contact weapon performs better at lower frequencies than the diffusible weapon. Snapshots show cropped (150μm) sections of the 300um-wide, 2-D simulation domain; below the black contour (arrow) represents the lethal concentration for the diffusible toxin. Scale bars: 50μm; innoculum: 100 cells. **D** Quantification of competition outcomes for two initial cells densities: ‘low’ (10 cells inoculum) and ‘high’ (100 cells inoculum). Competitive advantage assesses the log fold change in the attacker strain compared to its competitor from the beginning to end of the simulation (Methods). **E** Snapshots of competition outcomes for invasion scenario (invader frequency: 1%), with contact-dependent and diffusible toxin-armed attackers colored as in C; scale bars 10um. Successive timepoints (rows) show fates of initially rare attackers following random inoculation into confluent biofilms of susceptible cells. **F** Quantification of competition outcomes for invasions using the same competitive advantage metric as in D, quantified as a function of toxin secretion rate.

### Modelling predicts distinct strengths of short and long-range weapons

We use our model to compare short and long-range bacterial weapons that – other than their range of effect – are as similar as possible. Attacking cells can ‘fire’ a short-range weapon at random from their cell surface, intoxicating any susceptible cell(s) that are contacted (Figure 1A), which is intended to simulate contact weapons including type VI secretion systems^28,29^ and contact-dependent inhibition^13^. Alternatively, attacking cells can release a diffusible factor into the environment, which could represent range of toxins, including a small-molecule antibiotic, a ribosomally-synthesized bacteriocin, or a tailocin. The rate at which toxins are exported out of attacking cells is matched for short- and long-range weapons, and is controlled via a secretion rate parameter, *k*_*sec*_. Toxins also have matched potencies: both contact and diffusible toxins are lethal once intracellular concentrations exceed a set threshold *T*_*c*_. We assume that producing either toxin incurs an equivalent growth rate cost in attacking cells that is proportional to the secretion rate, i.e. increasing the toxin secretion rate results in a lower growth rate (Figure 1B) (Methods)^33,34^.

We begin by modelling competitions between an attacker strain, which either has a short- or a long-range weapon, and a second strain that is susceptible to the weapon. We look at a wide range of competition scenarios, varying the initial frequency of the attacker, initial density of cells, and the amount that the attacker invests in its weapon (i.e. toxin secretion rate). In each case the two strains are allowed to grow and interact for a set period of time (10h, Figure 1C), after which we compare the final attacker : susceptible cell ratio to its initial value (‘competitive advantage’, Methods). Both weapons tend to perform better when cells are seeded at high initial density, because this promotes cell-cell contact for short-range weapons^35^ and toxin accumulation for long-range weapons. Nevertheless, there remains a clear difference between the two weapon types. Across the majority of scenarios tested, the contact-dependent weapon provided an advantage to the attacker (Figures 1D, S1A). By contrast, the long-range weapon is more sensitive to starting conditions, requiring a higher secretion rate to be effective (Figure S1B), and only reaching its full potential when the attacker is seeded at high frequency and high density.

The models suggest a particular advantage of contact-dependent weapons: they are effective even when users are at a numerical disadvantage. This benefit is likely to be most significant when invading an established population, because here invaders will be outnumbered by residents. We explored this scenario further by modelling established biofilms of susceptible cells and simulating the late arrival of attacker cells (by replacing some of the susceptible cells at random with attackers). In this scenario, cells using the short-range weapon were able to successfully invade an established population, where increasing toxin secretion rate increased their success (Figures 1E, 1F, S1C)). Conversely, long-range weapons never enabled invasion, as producer cells could never amass in sufficient numbers to kill the susceptible strain.

In summary, even though we closely matched the properties of the two weapon types, the models predict that they perform very differently across the variety of competition scenarios we tested. In general, contact weapons work well across a range of frequencies, including cases when a strain is rare, allowing it to invade. By contrast, long-range weapons only function well when a producing strain is relatively abundant, but here they are an effective form of attack.

### Using genome editing to generate strains for weapon comparisons

Our modelling predicts that a long-range weapon will perform poorly at low attacker frequency, which is consistent with the findings of several previous studies – both theoretical and empirical – showing that toxin production is most effective when attackers are abundant^34,36–40^. However, to test our predictions on the relative benefits of short versus long-range weapons, a well-controlled comparison of the two types of weapons is required. To do this, we turned to the opportunistic pathogen *P. aeruginosa* (strain PAO1), which naturally carries both short- and long-range weapons. Of its long-range weapons, the mechanical tailocins (specifically its R pyocins) are known to be highly effective weapons under biofilm-like conditions^41,42^. As for short-range weapons, *P. aeruginosa* has a dedicated antibacterial type VI secretion system^16^, but it is under complex regulation and typically only fires in response to incoming attacks^28,43,44^. Therefore we instead chose to focus on a contact dependent inhibition (CDI) system of *P. aeruginosa*^45^ as a short-range weapon to test predictions.

For both weapons, we used genome editing to generate a strain that is susceptible to the weapon but otherwise well-matched to the attacker, allowing us to study the effects of each weapon on bacterial competition. For CDI, this was straightforward: deleting the three gene locus that encodes the CDI transporter, toxin filament and immunity (PA0040-PA0041), resulting in a strain that does not have CDI and is susceptible to CDI. For the tailocins, we selected pyocin R2, but here resistance is more complex as it is determined by the composition of lipopolysaccharide (LPS) moieties on the outer membrane^46^. Here, we engineered a susceptible strain by deleting both the pyocin R2 locus and the gene *wbpL*, which causes a deficiency in the LPS that in our strain background leads to susceptibility to pyocin R2 (Figure S2). However, this deficiency in the LPS caused a competitive disadvantage to the susceptible strain, independent of the effects of tailocins (Figure S3). To generate a near-matched attacker strain, therefore we made a Δ*wapR* deletion in the wildtype strain, which causes a similar LPS deficiency but not one that leads to pyocin R2 susceptibility (Figure S2, Methods).

In the absence of the effects of pyocin R2 (hereafter ‘tailocin’), the *wapR* mutation puts the attacker at a moderate disadvantage relative to the *wbpL* mutation in the susceptible strain (Figure S4). We quantified this difference and used it to adjust the predicted competitive advantage of the attacker strain in our experiments, in order to estimate the effects of the weapon alone (Figure S4, Methods). However, in practice this adjustment has little impact on the data because the benefits of the tailocin, when they are seen, massively outweigh this moderate cost of *wapR* deletion (for further discussion of these LPS mutations, see Methods).

### Experiments with P. aeruginosa shows distinct advantages to short- and long-range weapons

With these strains, we could then test our modelling predictions for a scenario where attackers armed with either a contact or diffusible weapon compete against a susceptible strain (Methods). We performed competitions when cells are growing on agar (the ‘colony biofilm model’)^47,48^, because this allows us to capture the dense, spatially-structured conditions thought to be typical of bacterial communities^49^, and because both weapons are expected to function well in this context (see Methods). The interior and edge of bacterial colonies represent distinct competition scenarios, due to the much greater potential for population expansion at the edge^50,51^. We therefore decided to sample each region separately (Methods), although in practice the competition outcomes are similar between the two. Consistent with previous work, we find that both systems have the potential to provide large competitive benefits for an attacking strain^42,45,52,53^. However, we also find that the outcome is strongly dependent upon the conditions of a given competition.

As the models predict, high attacker frequency is most important for the functioning of the long-range weapon (tailocin) (Figure 2B). In comparison, the short-range weapon (CDI) almost always performs equivalently or better at intermediate and low frequencies, especially at the colony edge (Figure 2C). This includes cases where the performance of CDI peaks at intermediate frequency as in the model (compare Figure 1D, high density, with Figure 2B, density 10^4^ cells). We also observe an improvement in the long-ranged weapon as initial cell densities increase. This pattern is again predicted by the model (Figure 1), although the effect is substantially stronger in the experiments where we can study much larger total numbers and ranges of initial cell density. Fluorescence micrographs of the colonies show how the competitions play out in space (Figure 2A, S5), and reveal differences between the middle and the edge of colonies, with the tailocin performing relatively poorly at the colony edge compared to CDI (Figure 2A middle bottom). This pattern is consistent again with the requirements for tailocins to build up to be effective, as this build up is expected to happen first in the colony interior and the colony edge last.

**Figure 2:**
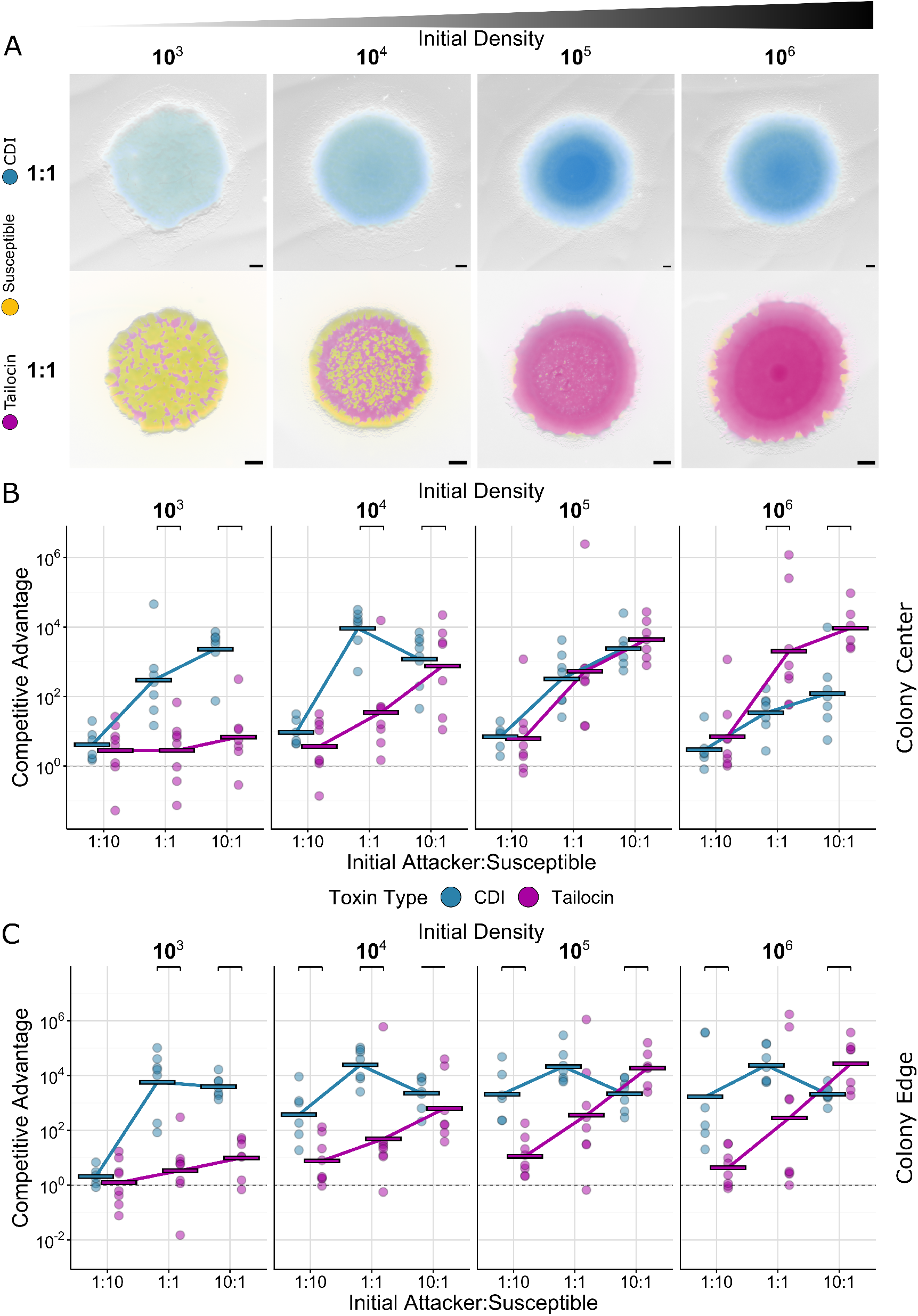
Experiments show the importance of high density and high frequency for long-range weapons. Colony competitions with *Pseudomonas aeruginosa* PAO1 between WT cells and mutants susceptible to either the contact dependent inhibition system (CDI), or pyocin R2 (Tailocin) inoculated from different densities (mean inoculum density 1.9 * 10^3^, 10^4^, 10^5^, 10^6^ CFU/μL). **A** Representative microscopy images from equal frequency (1:1) competitions after 48 h of growth. All strains express constitutive fluorescent protein genes and are false-coloured either blue (CDI attacker, top), magenta (tailocin attacker, bottom) or yellow (susceptible, top and bottom). Scale bar: 500 μm. **B** Quantification of competition outcomes at the colony center. **C** Quantification of colony competition outcomes at the colony edge. For B and C, competitions were assessed via counts of colony forming units. Competitive advantage assesses the log fold change in the attacker strain compared to its competitor from the beginning to end of the competition. The tailocin attacker advantage in B and C has been adjusted for a disadvantage in the background genotype of the attacking strain (see methods and Figures S3 and S4). Lines indicate the mean of replicates (n ≥ 6). Top brackets indicate a significant difference between the weapons (two-sided Welch’s t-test, p < 0.05, Benjamini-Hochberg MHT corrected 0.95).

In summary, our experiments show that both weapons can be highly effective, but that the two weapons perform best under different conditions. Tailocins are extremely effective at high frequencies and densities, while CDI performs more consistently across conditions, including the unique ability to provide a competitive advantage when a strain starts out rare and at low density.

### Head-to-head contests of short and long-range weapons

Our first experiments examine the performance of each weapon type against susceptible cells that do not fight back. We next explore the case where users of the two weapons meet. In this situation, a weapon can potentially take on new significance, as eliminating competitor cells also serves to reduce incoming attacks. The agent-based model again predicts that both initial frequency and cell density can be critical to the performance of the weapons (Figure 3AB, S6, Movie S2).

**Figure 3:**
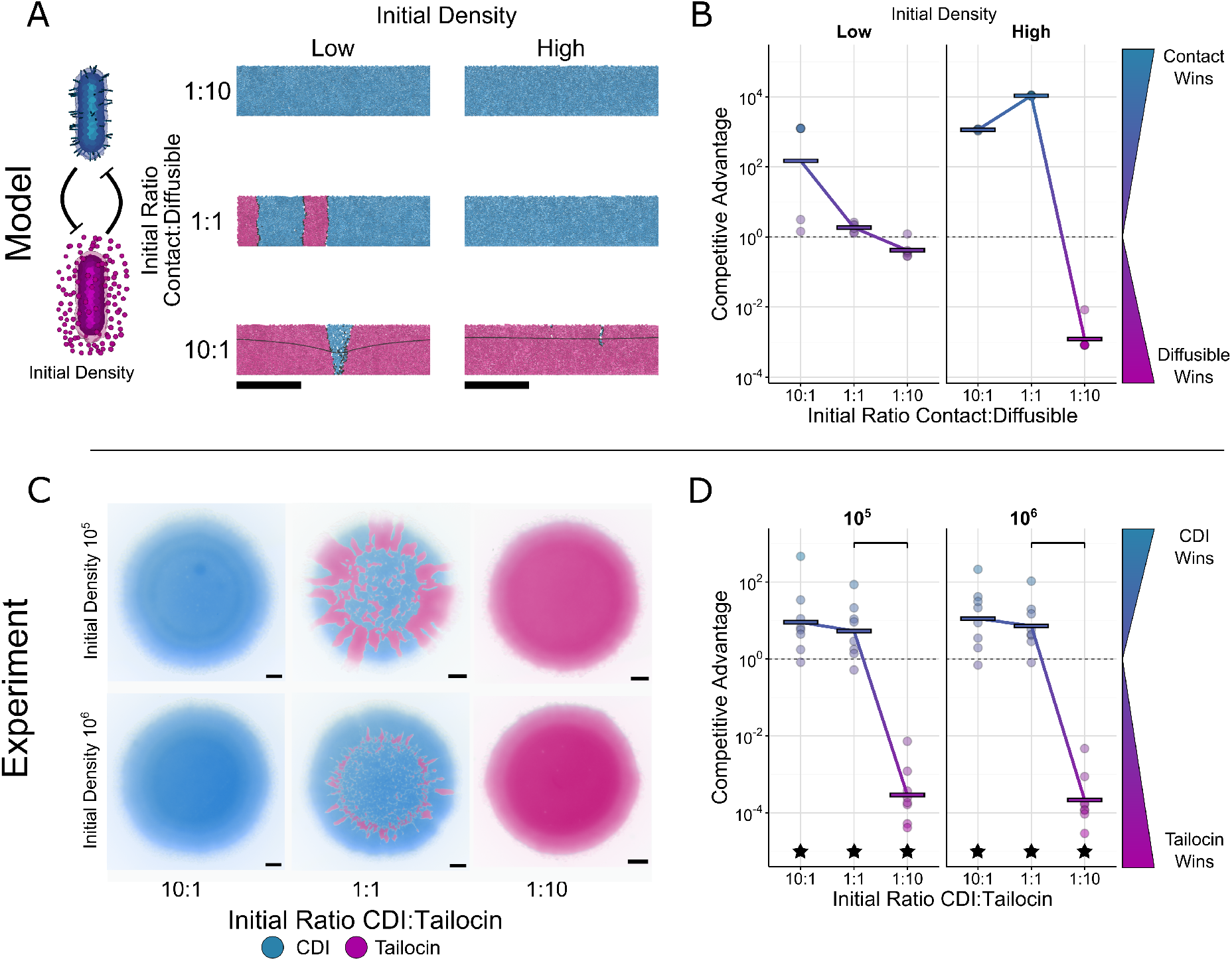
Head-to-head competitions between short and long-range weapon users. **A** Modelling: snapshots of competition outcomes for cells armed with contact-dependent toxins, but susceptible to diffusible toxins (blue cells) or cells armed with diffusible toxins, but susceptible to contact-dependent toxins (magenta cells). Both cells die upon lethal toxin exposure (black cells). Snapshots show cropped (150μm) sections of the 300um-wide, 2-D simulation domain; below the black contour (arrow) represents the lethal concentration for the diffusible toxin. Scale bars: 50μm; initial densities were ‘low’ (10 cells) or ‘high’ (100 cells). **B** Modelling: quantification of competition outcomes for two initial cell densities: ‘low’ (10 cells) and ‘high’ (100 cells). Competitive advantage assesses the log fold change in the attacker strain compared to its competitor from the beginning to end of the simulation (Methods). **C** Experiments: representative microscopy images of competitions between mutually susceptible CDI and tailocin producing cells after 48hrs inoculated from different densities (mean inoculum density 2.3 * 10^5^, 10^6^ CFU/μL). All strains are expressing constitutive fluorescent proteins and are false-coloured either blue (CDI attacker, tailocin susceptible) or magenta (tailocin attacker, CDI susceptible). Scale bar: 500 μm. **D** Experiments: quantification of colony competition outcomes via counts of colony forming units. Values above 1 (dashed line) indicate an advantage for CDI, while values below 1 indicate an advantage for tailocins. Data are adjusted to account for differences in competitiveness of the strain backgrounds (ΔwapR relative to ΔwbpL; see methods and Figure S2). Lines indicate the mean of replicates (n ≥ 6). Top brackets indicate a significant difference between the initial ratios (two-sided Welch’s t-test, p < 0.05, Benjamini-Hochberg MHT corrected 0.95). Stars indicate a significant competitive advantage (one-sided Welch’s t-test, p < 0.05, Benjamini-Hochberg MHT corrected 0.95). The genotype of the CDI using, tailocin susceptible strain (blue) is ΔR2ΔwbpL. The genotype of the tailocin using, CDI susceptible strain is ΔwapRΔCDI.

At low cell density, the impact of both weapons is limited, but the contact weapon user does gain an advantage when it starts in the majority. At high cell density, the long-range weapon user can win but, as in the single weapon competitions (Figure 1D), this requires it to start at high frequency. When the long-range strain is at low or equal initial frequency, the contact weapon performs the best.

To test these predictions, we competed a strain susceptible to CDI against a strain susceptible to tailocins (pyocin R2) (Figure 3CD). The CDI attacker was thus an LPS mutant (Δ*wbpL*, above, Methods), which renders it susceptible to the tailocin. As this mutation causes changes to the cell envelope, we first checked that the CDI system remained functional. The gene deletion (Δ*wbpL*) did not prevent CDI from functioning, but it did reduce the advantage provided by CDI at lower initial densities (Figure S7). Going forward, therefore, we only provide data from the two highest initial cell densities (10^6^ or 10^5^ cells/μL). We also focus on data from the interior of the colony going forward, as it is most reflective of typical biofilm growth ^32^, but data from the colony edge is similar (Figure S8). As predicted by the high cell-density model, only the CDI using strain is able to gain a significant advantage in equal-frequency competitions (Figures 3D). Moreover, again as predicted, both weapon users are able to win when they start in the majority. This effect is particularly strong for the tailocin, which provides a large competitive advantage when starting from high frequency.

### Contact and diffusible weapons are complementary

Our findings thus show that short- and long-range weapons can provide distinct advantages, which helps to explain why bacteria would carry both types. However, it is possible that these advantages are not provided simultaneously in practice. Therefore, we sought to experimentally study the effects of a single strain using both weapons, and compare it to each weapon being used alone. To allow susceptibility to tailocins and make the comparison as fair as possible, we put all attacker strains in the same LPS genetic background (Δ*wapR*), and all susceptible strains in the Δ*wbpL* background. All competition outcomes are again adjusted for the moderate disadvantage caused by the Δ*wapR* mutation, and we focus on high initial cell-densities as before (Figure S9, above). As predicted by the study of each weapon individually, across the competitions and contexts, the two weapons function in a complementary fashion, providing an equivalent or greater benefit than either weapon alone (Figure 4; Figure S9, S10). That is, we find that carrying both CDI and tailocins can allow a strain to receive benefits from both.

**Figure 4:**
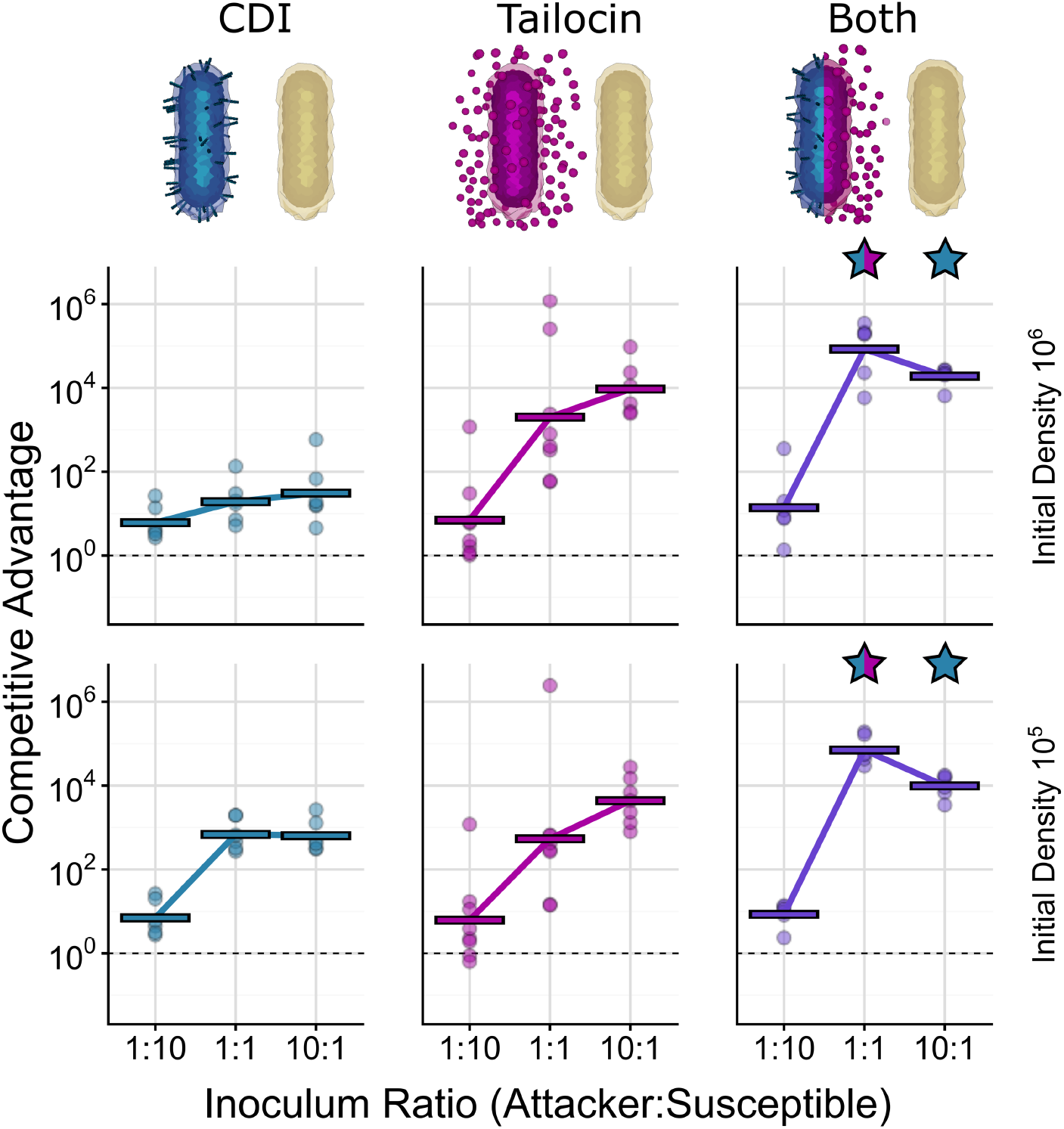
The benefits of short and long-range weapons combine positively in *P. aeruginosa*. Quantification of competition outcomes in the colony center for two initial cell densities (mean inoculum density 1.9 * 10^5^, 10^6^ CFU/μL). Competitive advantage assesses the log fold change in the attacker strain compared to its competitor from the beginning to end of the competition. Competitions where the attacker has just CDI (blue, left), just tailocins (magenta, centre) or both weapons (purple, right) show the advantage gained from using two weapons together as compared to just one. Data are adjusted to account for differences in competitiveness of the strain backgrounds (Δ*wapR* relative to Δ*wbpL*; see methods and Figure S2). Horizontal lines indicate the mean of replicates (n ≥ 6). Stars above the double weapon data indicate a significant difference between the combination of weapons and either single weapon (blue and magenta), or just CDI (blue) (two-sided Welch’s t-test, p < 0.05, Benjamini-Hochberg MHT corrected 0.95).

## Discussion

Bacteria use a vast variety of weapons to inhibit and kill competitors. Here we have shown that two major categories of weapons – short- and long-range – can provide distinct and complementary advantages to bacteria. Weapons that rely on contact between cells are generally effective whether a strain is rare or common. Conversely, weapons that diffuse across long ranges are more reliant on high producer cell frequency and density. However, under these conditions, long-range weapons can be particularly powerful in eliminating both armed and unarmed competitors. These observations help to answer two related questions. First, why do short- and long-range weapons exist at all in bacteria? Here, our work suggests that a strain may benefit more from one weapon type or another, depending on its ecology. For example, if bacterial fitness is more determined by the ability of a given strain to invade communities than persist in a community, contact-dependent weapons may be most useful. This matches with observations showing that pathogens including *Salmonella*^54^, *Shigella*^55^, and *V. cholera* ^56^ use the type VI secretion system (T6SS) during invasion of established communities, although gut-resident *Bacteroides fragilis* can also use its T6SS to repel invaders^57^ underlining the flexibility of short range weapons (Figures 1–3).

The second question is: why do many bacteria carry both short and long-range weapons? Again, the fact that the two weapon types can perform distinct functions provides an answer to why a cell would carry both. However, the possession of multiple weapons may also carry other non-mutually exclusive advantages. If two weapons are simply more potent than one, this can also favour carrying several of them^58^. In addition, if a given competitor is likely to carry resistance to some of the available weapons, natural selection may favour using multiple weapons simply to increase the likelihood that the competitor is susceptible to at least one. This explanation may be particularly important for cases where bacteria carry several of the same weapon type and functional differences are less pronounced e.g. *P. aeruginosa* releasing multiple S pyocin protein toxins and tailocins simultaneously via cell lysis^15^.

If the use of multiple weapons can be so beneficial, why is it not more widely seen in organisms other than bacteria? Short-range weapons, like CDI and T6SSs, can be used against competitors by lone cells. The effectiveness of long-range weapons, meanwhile, rests upon the strength in numbers that comes with group living. There are some examples of group combat strategies in animals. Colonial marine invertebrates engage in toxin-mediated competition^59^, and the Asian honeybee *Apis cerana* kills the giant Asian hornet *Vespa mandarinia* by ‘heat balling’, whereby the bees cover the hornet and vibrate their flight muscles to overheat it^60^. However, this latter example is best described as predator defense and, in general, group-level strategies appear to be the exception in most animal conflicts. Instead, most animal contests are dyadic, meaning they occur between just two individuals^61,62^. By contrast, mounting evidence suggests that bacteria commonly live, and fight, as part of large groups. This lifestyle gives them the opportunity to engage both as individuals and as collectives. Our work suggests that it is this propensity for group living that has led to the widespread evolution of both short- and long-range weapons.

## Methods

### Agent-based modelling

A major challenge in comparing bacterial weapons is that even within the broad categories of contact and diffusible, their method of deployment varies substantially. The type IV secretion system (T6SS) can be fired at random intervals to deliver toxins *or* only in response to incoming attack^43^, whereas CDI filaments are produced to decorate attacking cells in unknown numbers, but toxin translocation only occurs in response to receptor binding on a target cell. The small molecule antibiotics of *Streptomyces* are secreted from intact cells, whereas *E. coli*’s colicins and the pyocins of *P. aeruginosa* require cell lysis for release. These examples also demonstrate the high variability in the cost of using weapons. The use of computational models is very amenable to these challenges, as we can unify the production of both contact and diffusible weapons under a single parameter ‘secretion rate’ and make their growth costs (per unit secretion) equal.

The behaviours of cellular groups are often emergent, in the sense that they can only be understood in terms of collective effects that arise whenever there are many interacting organisms^63^. A strength of our modelling approach is that it is able to capture emergent and spatial effects, such as the importance of cell shape for bacterial competition^30,32^, the importance of lytic toxins for T6SS attack ^29^, and the need for strong reciprocation when the T6SS is used in response to incoming attacks^29^. For each of these examples, the models predicted novel biology that was subsequently validated empirically using experimental work^28,29,32^, giving us confidence in our modelling approach.

All simulations were carried out using CellModeller^31^, an open-source software platform for running agent-based models of bacterial growth. To model contact-based and diffusible toxin secretion, we implement additional python-based Cellmodeller modules, whose source codes are available here (<u>Smith Github</u>). The key processes incorporated in our model are summarised below; model variables and parameters are summarised in Tables S1 and S2, respectively.

#### Model description

##### Cell growth and division

In our simulations, bacterial cells are represented as short capsules with a fixed radius *R*=0.5μm, and a birth segment length of 0.01μm (equivalent to a birth volume *V*_*0*_ = 0.54 μm^3^). Cells grow through elongation, dividing after doubling their birth volume *V*_*0*_, plus a small random noise term *ξ*_*V*_ ~ *U(0, 0.05)V*_*0*_. Each simulation timestep *Δt*, each cell grows by an amount proportional to that cell’s current volume, *V’* = *k*_*grow*_*V*, discretised as *V(t+Δt)* = *V(t) (1*+*k*_*grow*_.*Δt)*. Here, *k*_*grow*_ [s^−1^] represents the net per capita growth rate. The production of ranged or contact toxins is assumed to be costly, such that weapon users suffer a growth penalty proportional to their toxin secretion rate, *k*_*grow*_ = *k*_*max*_*(1–ck_sec_)*, with *c*, k_max_, and *k*_*sec*_ the pro rata toxin cost, maximum growth rate and equivalent contact toxin secretion rate, respectively. Throughout, we assume that nutrient access is non-limiting, such that *k*_*max*_ is independent of cells’ positioning in a community.

##### Mechanical interactions

Mechanically, cells are modelled as rigid, elastic particles that push on one another as they grow and divide. Each simulation timestep, immediately following the growth stage outlined above, an energy penalty method is used to compute cell movements necessary to minimise total cell-cell overlap, subject to viscous drag forces acting on each cell. This process, described previously in detail^31,32,64^, approximates the elastic repulsion forces acting between cells in physical contact.

##### Contact-dependent toxins

As in previous publications^28,29^, we use a custom Python module to represent cell-cell antagonism via contact-dependent toxins. While this module was previously used to study T6SS-mediated interference, it is a generic representation of contact-dependent warfare that allows us to make general predictions that should apply to many mechanisms, including contact-dependent inhibition. Each simulation timestep, cells armed with contact toxins fire needles of length *R,* projecting orthogonally from randomly-chosen sites on their cell surface. The number of secretion events per cell per unit time is drawn from a Poisson distribution, whose mean is the secretion rate *k*_*sec*_. After firing, each needle is checked to determine if it comes into contact with any other cell in the population (line-segment method). Successful hits are logged for each target cell, and result in cell death if their number (excluding hits by kin cells) exceeds a lethal threshold *N*_*hits*_ = 1 ^65^.

##### Diffusible toxins

We assume toxins to be freely-diffusible solutes that kill susceptible cells when their local concentration *u*_*T*_ exceeds a lethal threshold *T*_*C*_ (controlled by *N*_*hits*_; see below). To represent natural variability in toxin susceptibility^66–68^, lethal toxin threshold is drawn for each cell from a normal distribution, *N(1, 0.2)* at birth (we ignore this stochasticity in the discrete contact toxin model, since with mean *N*_*hits*_ = 1, the chances of any cell surviving more than one hit are approximately 1:130,000). To model the toxin concentration field *u_T_ = u_T_(x, y*) [kg_T_m^−3^] for a given cell configuration, we use the reaction-diffusion equation *∂u_T_/∂t = D_T_∇^2^u_T_ + k_T_ɑ⍴ɸ(x, y)* ^69^. Here, *D_T_, k_T_, ɑ, ⍴,* and *ɸ(x, y)* are respectively the toxin diffusivity [m^−3^], the specific toxin production rate [s^−1^], the toxin yield per unit cell biomass [kg_T_kg_X-1_], the cell biomass density [kg_x_m^−3^], and the cell volume fraction function [unitless]. In non-dimensional form, pseudo-steady-state solutions to this equation are given by *∇^2^u_T_ = ⅅ_T_ ɸ(x,y)*. The behaviour of this equation is governed by a single parameter grouping, the Damköhler number *ⅅ_T_ = l^2^k_T_ɑ⍴ / D_T_T_c_*. Notably, with this scaling, increasing toxin production *k*_*T*_ or proportionally decreasing its diffusivity *D*_*T*_ or lethal concentration *T*_*c*_ are all equivalent, and so by varying *ⅅ*_*T*_ one can explore variation with respect all three of these parameters. We compute pseudo-steady-state solutions with the finite element method, using the FEniCS python library and supporting CellModeller modules. Solutions are evaluated on a 2-D rectangular domain of dimensions *L*_*x*_ by *L*_*y*_ with crossed mesh element size *h* = 5μm. Solutions are subject to mixed boundary conditions: periodic boundary conditions (left and right edges), Neumann boundary conditions (base edge), and Dirichlet boundary conditions (*u*_*T*_ = 0 along top edge). As in previous studies^69^, we assume that toxin is not subject to degradation or removal as part of its activity; toxin is only lost from the domain via leakage along the top edge.

##### Parity between diffusible and contact dependent toxins

Our model aims to compare diffusible and contact weapons in a like-for-like manner. For this comparison to be as fair as possible, we assume in all cases that both weapons involve the secretion of the same (hypothetical) toxin, at the same rate *k*_*T,cell*_, with the same potency (lethal concentration) *T*_*c*_, and at the same growth cost *c*. For simplicity, the secretion rate is fixed within each simulation, while in reality many bacteria responsively upregulate toxin production via competition sensing^70,71^ and other mechanisms^72^. Here we outline the equations that link these properties between the two models, bridging the (discrete) contact model with the (continuum) diffusible toxin model. For contact weapons, the per-cell secretion rate *k*_*T,cell*_ is given by *k_T,cell_ = k_sec_T_sec_*, with *k*_*sec*_ [h^−1^] the per-cell secretion rate, and *T*_*sec*_ [kg_T_] the mass of toxin released by each secretion event. For diffusible toxins, this can be expressed as *k_T,cell_ = k_T_ɑ⍴l*^*3*^ (terms defined as above) with *l*^*3*^ approximating the cell volume. The contact-dependent secretion rate *k*_*sec*_ can therefore be related to the diffusible toxin Damköhler number, *ⅅ*_*T*_, as *ⅅ_T_ = T_sec_k_sec_ / D_T_T_c_l.* To relate the potencies of contact and diffusible toxins, we assume that *N*_*hits*_ is equivalent to the minimum number of contact events required to raise the intracellular toxin concentration to the lethal threshold *T*_*c*_: *N_hits_ = T_c_l^3^ / T_sec_*, with *l*^*3*^ approximating a cell’s volume as before. Moreover, by combining these equations, we can eliminate the (unknown) contact toxin load *T*_*sec*_ and compute *ⅅ_T_ = k_sec_ (l^2^ / N_hits_D_T_)* for the two weapons operating at equivalent secretion rates. This relation highlights that toxin diffusivity *D*_*T*_ is a crucial parameter for our model, since increasing *D*_*T*_ is equivalent to reducing effective diffusible toxin production while keeping contact toxin production the same. While previous work has explored the interplay between toxin diffusivity, cost, and environmental structure^69^, here we assume a fixed value for *D*_*T*_ (that for colicin Ia^73^), which is towards the lower limit of the range explored by previous work^69^.

##### Simulation domain and protocols

All model simulations are run on a *L*_*x*_-by-*L*_*y*_ rectangular domain with lateral periodic boundary conditions, representing a vertical 2-D slice through a bacterial community. The base of the domain, *y* = 0, is an impenetrable substrate. The domain width *L*_*x*_ is fixed; its height *L*_*y*_ is set to track the maximum height *p*_*y*_ of bacterial cell groups as *L_y_ = max(p_y_) + δ*, with *δ* the diffusive boundary layer thickness. To represent cell detachment through mechanical sloughing, cells are removed from the simulation once they reach a height *h*_*slougher*_. All simulations are initiated by randomly scattering cells along the base edge of the simulation domain. Unless otherwise indicated, simulations run for a fixed duration of 400 steps (10h of simulated time, timestep *Δt* = 0.025h). Invasion simulations involve a 2-step process: first, the domain is inoculated as above using only susceptible cells, which are then allowed to divide until the cell group reaches the slougher height *h*_*slougher*_. Then, a random subpopulation of the susceptible cells is converted to the invading cell type, and the simulation is allowed to run for up to 1,000 steps (25h), terminating early if either cell type is lost from the simulation.

#### Computation and Postprocessing

All agent-based simulations were run using a 2017 Apple MacBook Pro laptop computer, with concurrent simulations distributed between an Intel 3.1 GHz quadcore i7-7920HQ CPU, an Intel HD 630 Graphics card, and an AMD Radeon Pro 560 Compute Engine. Simulation data were visualised using Paraview software, and analysed using custom Matlab and R scripts.

### Experiments

#### Strain Construction

Deletion mutants were constructed using standard two-step allelic exchange methods using the vector pEXG2 and Gibson assembly^74,75^. Primer sequences for up/downstream regions and exterior confirmation primers are listed in Table S3. Constructed deletion vectors were introduced into *P. aeruginosa* PAO1 by conjugation with *E. coli* JKE201^76^ and transconjugants were selected on LB agar with 50 μg/mL gentamycin. After counter-selection on LB no salt, 10% sucrose, colony PCR positive clones were confirmed by Sanger sequencing (Source Bioscience, Nottingham, UK) and gentamycin sensitivity confirmed. Stocks in LB 20% glycerol were stored at −80C. For strains with multiple deletions, they are listed in the order the deletions were made. Strains were subsequently constitutively tagged with eYFP and mScarlet (Sujatha Subramoni, unpublished) using pUC18-mini-Tn7-GmR ^77^ delivered by conjugation with *E. coli* S17λ and selected on Pseudomonas Isolation Agar with 100 μg/mL gentamycin. Replicates were performed with each attacker/susceptible combination carrying opposite fluorescent markers.

#### Engineering strains for weapon comparisons

For CDI mediated competition, we deleted the three gene locus that encodes the CDI transporter, toxin filament and immunity (PA0040-PA0041), which resulted in a strain that does not have CDI and is susceptible to CDI. For pyocin R2, we found that deletion of *wbpL* led to susceptibility. *wbpL* is a transferase which initiates the formation of LPS chains by adding the first sugar to the undecaprenol-phosphate carrier, and in a previous study a cosmid library-derived insertional inactivation mutant was found to be resistant to pyocin R2^46^. For our purposes, we made an in-frame deletion mutant of *wbpL* from scratch, which grew poorly in liquid media. We reasoned that this was due to inhibition by its own pyocins. Consistent with this, making the *wbpL* mutant in a pyocin R2 deletion background restored growth. Moreover, this strain was then found to be susceptible to pyocin R2 from its parent. Finally, to confirm that *wbpL* was responsible for this susceptibility we complemented the in-frame deletion mutant of *wbpL* with a copy of *wbpL* on a plasmid, which restored pyocin resistance (Figure S4). On this basis, we are confident that, in our strain background, that deleting *wbpL* leads to pyocin susceptibility.

*wbpL* was PCR amplified from *P. aeruginosa* PAO1 genomic DNA and inserted into the expression vector pSEVA524 (Rubén de Dios, Eduardo Santero and Francisca Reyes-Ramírez, unpublished)^78^ by Gibson assembly (NEB Hifi Assembly, NEB Location). After conjugation with *E. coli* JKE201 and selection on LB 10 ug/mL tetracycline a positive clone and a clone carrying the empty vector were tested for susceptibility to pyocin R2. Strains were grown overnight at 37°C in LB (with 10 ug/mL tetracycline), then 1 mL mixed with 7mL 0.75% LB agar and poured onto a warm LB plate to generate an overlay. Attacker cultures were also grown similarly and R pyocins were prepared by filter sterilizing supernatant from overnight cultures. 20 μL was spotted on the overlays which were then dried and incubated overnight at 37°C prior to photographing using a Gel Doc imager.

We also made a deletion of *wapR* which attaches the first L-rhamnose to the LPS core, initiating LPS capping, which did retain resistance to pyocin R2. We then had two strains Δ*wbpL* and Δ*wapR* that are LPS defective, but one is susceptible to pyocin R2 and one is not. These strains were thus used for the long-range diffusible weapon experiments.

#### Culturing and the colony biofilm model

Strains were recovered from cryo stocks by streaking on LB 1.5% agar and incubated overnight at 30°C. LB 1.5% agar for competitions was prepared immediately prior to competition setup by pouring 20 mL into a petri dish and allowing it to set for 15 minutes in a laminar flow hood. Colony competitions were prepared as previously by scraping cells off the overnight plate and resuspending cells to an initial OD600 of 1 ^47^. Strains were mixed at defined ratios of 1:10, 1:1 and 10:1 then serially diluted 10-fold and 1 uL spotted on the prepared plate to generate competitions at various initial ratios and densities. Initial culture density was determined by serially diluting and spot plating.

We performed competitions when cells are growing on agar (the ‘colony biofilm model’)^47,48^. In nature, most bacteria live in densely-packed communities, such as surface-associated biofilms, that are densely packed and spatially structured^49,63,79^. These high-density conditions are where both short and long-range weapons are expected to function at their best, because they ensure plentiful cell-cell contacts and the potential for factors released in the cells to build up to high concentrations^39,40,80,81^. Bacterial weapons can also strongly influence the spatial structure of competing strains, and vice versa; effects that are not captured in liquid culture^27,63,82,83^. Finally, *P. aeruginosa*’s CDI systems is upregulated in structured static cultures, as compared to shaking culture, again suggesting these are the conditions where it has most impact^45^. Growing bacteria on agar captures these dense and structured conditions in a highly tractable manner, allowing large numbers of competitions and conditions to be studied and imaged^32,47,84^ (Figure 2). Colonies also represents a good match to our model framework, which is again focused on high density, biofilm-like conditions.

#### Imaging of Colonies and Quantification of Competition Outcomes

After 48 h of growth at room temperature, colonies were imaged using a Zeiss Axio Zoom V16 microscope with a Zeiss MRm camera, 0.5X PlanApo Z air objective and HXP 200C fluorescence light source. 48 h allowed colonies inoculated from the lowest initial densities to grow sufficiently to be observed. All colonies from a single set of frequencies and densities were imaged at the same zoom (between 1-2.5x). To make the composite images shown in the figures, the display histograms of each channel were scaled to the minimum and maximum values found in the entire set of frequency and density, meaning images can be compared within sets (i.e. weapon competitions) but not between. For Figure S5, all images were treated as a set, so comparisons can be made across all images. After microscopy imaging, colonies were sampled with a 10 μL pipette at both the center and edge of the colony into 0.9% saline. Samples were homogenized, serially diluted and 5 μL spotted onto LB or LB 50 μg/mL gentamycin and incubated at 30°C overnight. Colonies were counted to determine the final ratio of the two strains.

#### Calculation of Competitive Advantage

Using the initial density counted from the original inoculum cultures and the known inoculum ratios, the initial ratio of attacker:susceptible strains was determined. The final ratio was determined from the CFU counts of the serially diluted center and edge samples, with a detection limit of 2000 CFU/mL used to replace zeros and prevent dividing by zero. Competitive advantage was defined as the log of the final ratio / initial ratio.

## Author Contributions Statement

SCB, WPJS and KRF designed the study, analyzed the data, wrote and edited the manuscript. SCB designed and performed the experiments. WPJS implemented and performed the modelling. WPJS, SCB and KRF designed and interpreted the models.

## Acknowledgements

The authors would like to think OPC for the pEXG2 vector and JKE201 conjugation strain, SS for the pUC19-Tn7-mScarlet plasmid, and RD, ES, FRR and for the pSEVA524 complementation vector. We are indebted to Carey Nadell and Oliver Meacock for comments and feedback during manuscript drafting. This project is supported by a Templeton World Charity Foundation grant. KRF and WPJS are supported by European Research Council Grant 787932, and by Wellcome Trust Investigator award 209397/Z/17/Z. WPJS is also funded by a Sir Henry Wellcome Postdoctoral fellowship award, 222795/Z/21/Z.

