## SupplementaryInformation for "Bows and swords: why bacteria carry short and long-range weapons"

**Supplementary tables**

Table S1: Model Variables

| Variable | Symbol | Units |
|---|---|---|
| Cartesian coordinates | $x, y$ | μm |
| Cell volume fraction field | $\phi(x,y)$ | - |
| Toxin concentration field | $u_T(x,y)$ | $kg_T m^{-3}$ |
| For each cell i: | | |
| Position vector | $\boldsymbol{p_i} = (p_x, p_y, p_z)_i$ | μm |
| Orientation unit vector | $\boldsymbol{a_i} = (a_x, a_y, a_z)_i$ | - |
| Segment length | $L_i$ | μm |
| Volume | $V_i = 4\pi R^3/3 + \pi L_i R^2$ | $\mu m^3$ |
| Specific growth rate | $k_{max}(1–ck_{sec})$ | $h^{-1}$ |

Table S2: Model Parameters

| Type | Parameter | Symbol / equation(s) | Value(s) [units] | Source |
|---|---|---|---|---|
| * No specific value assumed; parameter varied implicitly via cluster $\mathbb{D}_T$. ‡ Controlled by $N_{hits}$. | | | | |
| Cells | Cell radius | $R = l/2$ | 0.5 [μm] | [1] |
| | Cell volume at birth | $V_0$ | 0.54 [$\mu m^3$] | [2] |
| | Max cell growth rate | $k_{max}$ | 1.0 [$h^{-1}$] | [1] |
| | Cell biomass density | $\rho$ | * [$kg_X\ m^{-3}$] | - |
| | Random noise in division volume | $\eta_{div}$ | 5.0 [%] | [2] |
| | Cell division orientation noise | $\eta_{orient}$ | 0.2 [%] | [2] |
| Toxin -Common | Per cell secretion rate | $k_{T,cell}$ | * [$kg_T\ s^{-1}$] | - |
| | Lethal concentration | $T_c$ | * ‡ [$kg_T m^{-3}$] | - |
| | Lysis delay | $1 / k_{lysis}$ | 0.125 [h] | [3] |
| | Weapon cost per unit secretion | $c$ | 0.0005 [h] | This study |
| Toxin - Contact | Extracellular needle length | $L_{needle} = R$ | 0.5μm | [3] |
| | Min. needle penetration for hit | $L_{penetration}$ | 0.01μm | [3] |
| | Number of hits to kill target cell | $N_{hits}$ | 1 [-] | Estimated from [4] |
| | Secretion rate | $k_{sec}$ | 0.0-500.0 [$h^{-1}$] | This study |

| | | | | |
|---|---|---|---|---|
| Toxin - Diffusible | Toxin specific production rate | $k_T$ | * [s] | - |
| | Toxin yield per unit biomass | $\alpha$ | * [$kg_T\ kg_X^{-1}$] | - |
| | Toxin diffusivity | $D_T$ | $4x10^{-11}$ [$m^2s^{-1}$] | [5] |
| | Toxin Damköhler number | $\mathbb{D}_T = l^2 k_T \alpha \rho / D_T T_c$ $= k_{sec}\, l^3 / D_T\, N_{hits}$ | $0\text{-}3.5x10^{-3}$ [-] | This study |
| Domain | Diffusive boundary layer height | $\delta$ | 25 [μm] | This study |
| | Domain width | $L_x$ | 300 c | This study |
| | Slougher height | $h_{slougher}$ | 40 [μm] | This study |
| Numerical | Mesh element size | $h$ | 5 [μm] | [2] |
| | Simulation timestep | $\Delta t$ | 0.025 [h] | [1] |
| | Cell / needle sorting grid size | $h$ | 10 [μm] | [3] |
| | Conjugate gradient absolute tolerance | $e_{CG}$ | 0.001 [-] | [1] |
| | Max. contact iterations | $Max_{iter}$ | 8 [-] | [1] |
| | Regularization weight | $\alpha$ | 0.04 [-] | [2] |
| | Growth restriction factor | $1/\gamma$ | 0.002 [-] | [2] |

Supplementary Table 3: Primers used for construction of deletion mutants and *wbpL* complementation.

| Primer | Sequence |
|---|---|
| CDI1-out-F | agttcatgtccaatcaccacacc |
| CDI1-out-R | agggttcagttcgatgtaccgg |
| CDI1-del-UpF | aagcttctgcaggtcgactctagaggatccaagaagatcgaactggtcggc |
| CDI1-del-UpR | tgtacttcggcataatcagcctcctccagcgcgatggagagccacgatttc |
| CDI1-del-DownF | tcaaggaatgaaatcgtggctctccatcgctatatagttgagtaagctttgctgcga |
| CDI1-del-DownR | cccgtggaaattaattaaggtaccgaattcaagcgattccgatatcgtgtcg |
| R2-out-F | gaggctttccatggctgacc |
| R2-out-R | tgaggttgcaggtgacatcc |
| R2-del-UpF | aagcttctgcaggtcgactctagaggatccgatcacgccaacgaactggtc |
| R2-del-UpR | gatgggtttcaggcgtcaccccttgccgccagcctgttcaggcatggttg |
| R2-del-DownF | tcgaaggagtcaaccatgcctgaacaggctggcggcaaggggtgac |
| R2-del-DownR | cccgtggaaattaattaaggtaccgaattctacgttgtaccgaccgattccc |
| wapR-detect-F | ccatgtctatggcgccttcac |
| wapR-detect-R | aaccgcatccgtttcttcgtc |
| wapR-del-DownF | caagcttctgcaggtcgactctagaggatcgctactgggctacatcctgtac |
| WapR-del-DownR | taaggttgtagttatggcgctcgtcgatgagaagtactggaacgagaagaccttc |
| wapR-del-UpF | ccgcggaaggtcttctcgttccagtacttctcatcgacgagcgccataac |
| wapR-del-UpR | acccgtggaaattaattaaggtaccgaattcaagaatttcgaggcgatggc |
| wbpL-detect-F | ggctcagtatagccggtaagtc |
| wbpL-detect-R | caaatacgaggctgagcaggag |
| wbpL-del-DownF | caagcttctgcaggtcgactctagaggatccaggcggtgatagccaaagtg |
| wbpL-del-DownR | aaggttctctttccaatgatgatctggatgctcttggcggtaggatacaagg |
| wbpL-del-UpF | ggaacccgccttgtatcctaccgccaagagcatccagatcatcattggaaagagaac |
| wbpL-del-UpR | ACCCGTGGAAATTAATTAAGGTACCGAATTccgcctttgatctatgccaatg |
| wbpL-compF | tcggtacccgggctagatttaagaaggagatatacatatgatgatctggatgatcgcgtg |
| wbpL-compR | gtcgccagggttttcccagtcacgacgcggccgcattaggatttttccaaggaacccgc |

### Supplemental Figures

### Captions for supplementary videos

**Movie S1: Comparison of short and long-range weapons using agent-based modelling.** Both attacker types compete against susceptible cells but the long-range (diffusing) toxins are only effective a high initial frequency (bottom row). By contrast, the short-range (contact) weapon is effective at all frequencies. Simulated cells compete for space within a 300μm wide, 2D-biofilm. Models are initiated with either 100 total cells on the surface, and with an initial ratio of armed attackers (contact weapon, blue; diffusible weapon, magenta) to unarmed susceptibles (yellow) of 1:10 (top), 1:1 (middle) or 10:1 (bottom). Simulations last for 10.025 h.

**Movie S2: Agent-based modelling of direct contests between the two weapons users.** Here contact and diffusible weapon compete against each other. When starting at higher frequency, both can overcome the other, but at equal frequency it is the contact weapon user that wins. Simulated cells compete for space within a 300μm wide, 2D-biofilm. Models are initiated with 100 total cells on the surface, and with an initial ratio of contact weapon armed cells (blue) to diffusible weapon armed cells (magenta) 10:1 (top), 1:1 (middle) or 1:10 (bottom). Simulations last for 10.025 h.

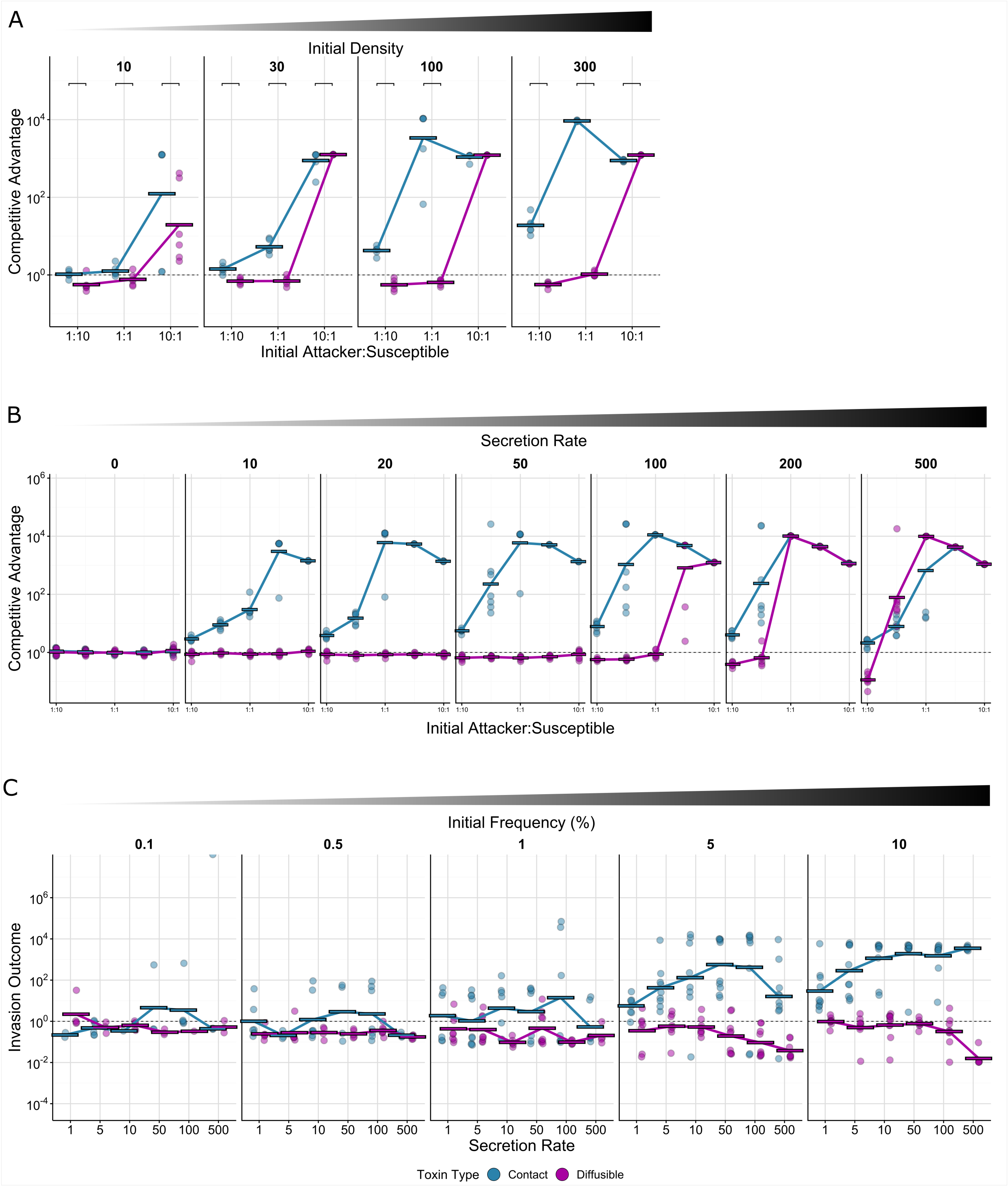


**Figure S1: Agent-based modelling shows differences between weapons due to initial density of competitions, weapons depend differently on toxin secretion rate and that contact weapons better facilitate invasion than diffusible weapons at equivalent secretion rates. A** Quantification of competition outcomes for all tested densities (secretion rate: 100). Densities of 10 and 100 cells correspond respectively to “Low” and “High” starting densities shown in Figure 1. Competitive advantage assesses the log fold change in the attacker strain compared to its competitor from the beginning to end of the simulation (Methods). **B** Outcomes of competition simulations over a range of secretion rates and initial attacker frequencies (Initial density: 150 cells). Competitive advantage assesses the log fold change in the attacker strain compared to its competitor from the beginning to end of the simulation (Methods). **C** Quantification of competition outcomes for invasions as a function of secretion rate. Invasion outcome is the same as competitive advantage (the log fold change in the attacker strain compared to its competitor from the time of invasion to the end of the simulation).

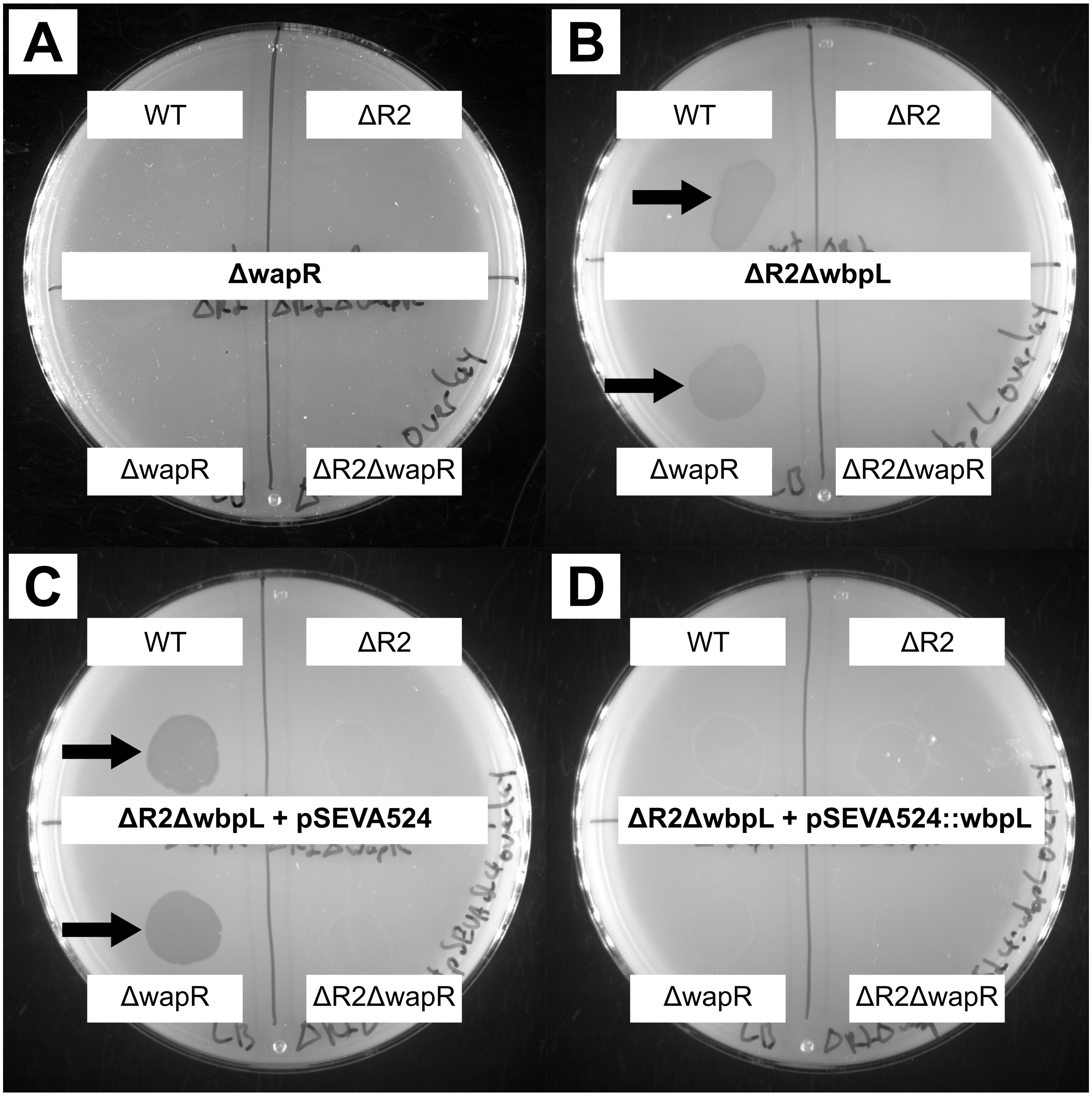


**Figure S2**: **Lipopolysaccharide biosynthesis genes affect susceptibility to pyocin R2.** Images of agar overlay assays showing pyocin R2 zones of clearing (arrows) for different LPS mutants. The strain in the overlay is indicated in the center of each plate. The source strain for the pyocin R2 is indicated at the corners. Pyocins were prepared by sterile filtering supernatant from overnight cultures. Overlays were prepared by mixing 1mL of overnight culture with 7mL 0.75% LB agar then thoroughly drying. **A** Δ*wapR* shows no zones of clearing. **B** ΔR2 Δ*wbpL* (the entire pyocin R2 gene cassette was first deleted from this strain, then *wbpL* deleted second) shows zones of clearing from WT and Δ*wapR*, but not when pyocin R2 is deleted. **C** Complementing ΔR2 Δ*wbpL* with empty vector pSEVA-524 does not rescue clearing. **D** Complementing ΔR2Δ*wbpL* with pSEVA-524 carrying *wbpL* shows no clearing from WT or Δ*wapR*.

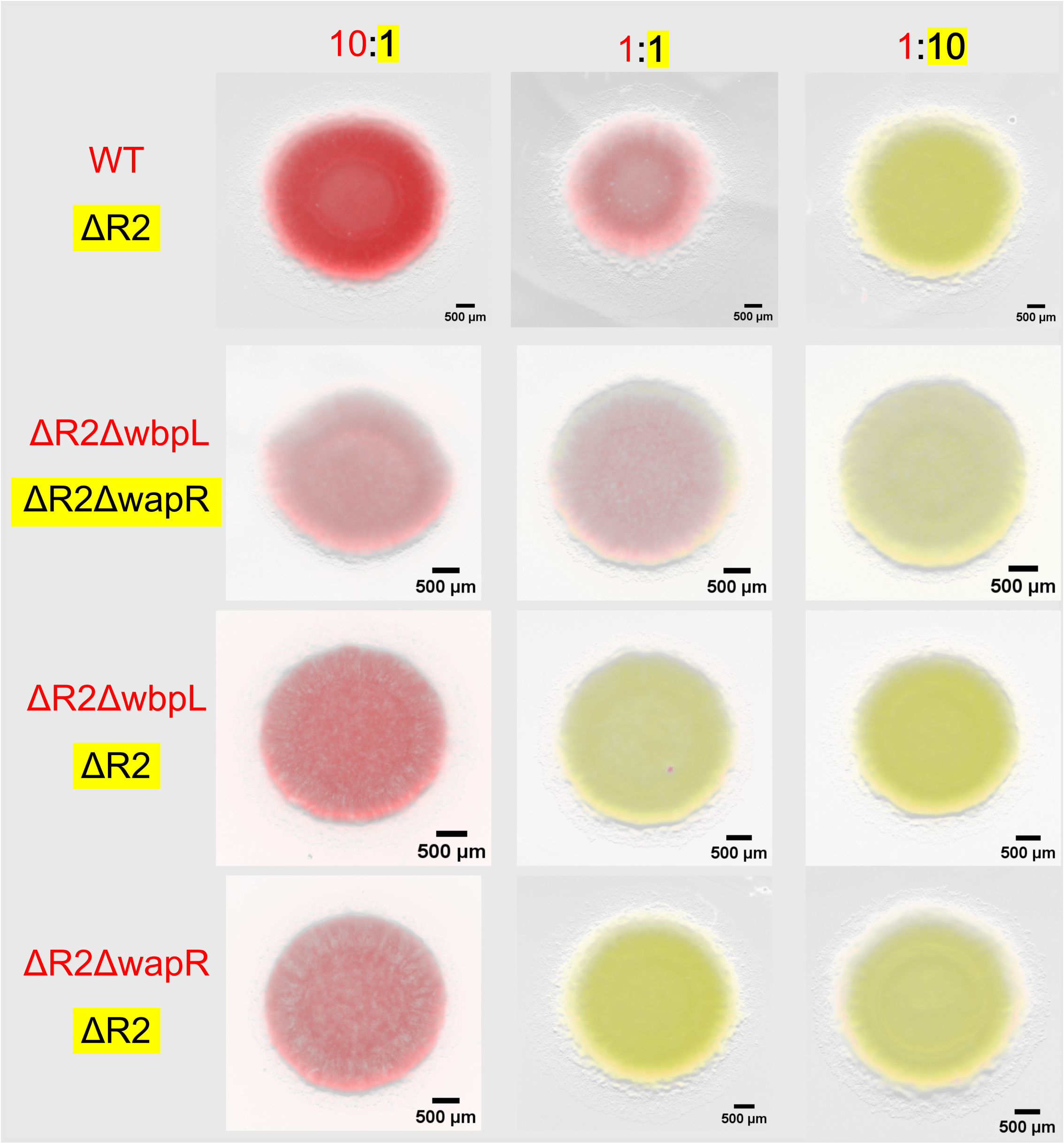


**Figure S3: Lipopolysaccharide biosynthesis gene deletions affect competition outcomes in the absence of pyocin R2.** Microscopy shows that the Δ*wbpL* strain cannot compete with wild-type *P. aeruginosa*, even in the absence of killing by tailocins (pyocin R2). Conversely, Δ*wbpL* and Δ*wapR* (second from top) are closely matched and look similar to wild-type competed against ΔR2 (top). Competitions were inoculated with ~2x$10^6$ cells/µL.

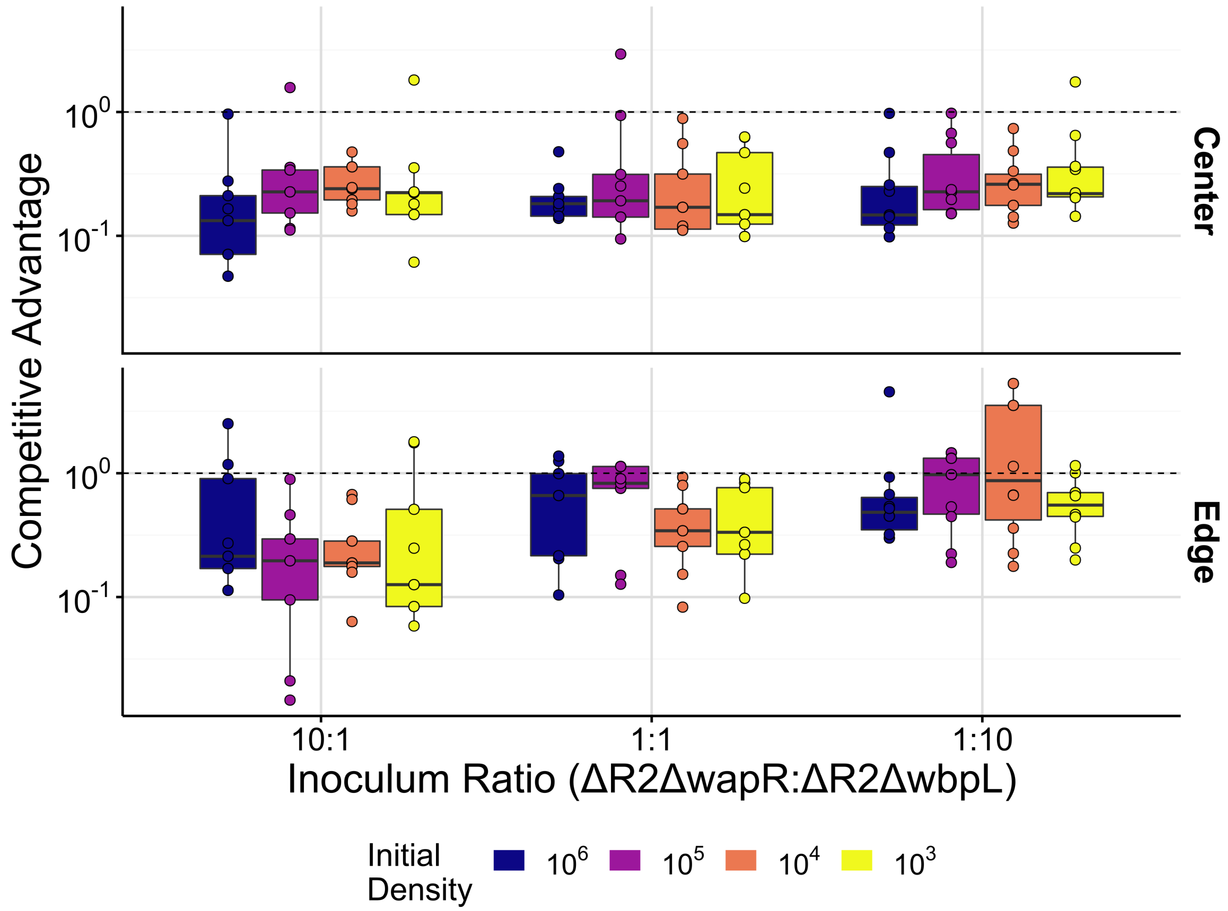


**Figure S4: Deletion of lipopolysaccharide biosynthesis gene *wapR* causes a disadvantage compared to deletion of *wbpL* when both strains have pyocin R2 deleted.** Quantification of colony competition outcomes between lipopolysaccharide biosynthesis mutants in the absence of pyocin R2. Colonies were inoculated at the stated initial ratios and densities (mean inoculum density 1.8 * $10^3$, $10^4$, $10^5$, $10^6$ CFU/µL). Competitive advantage assesses the log fold change in the attacker strain compared to its competitor from the beginning to end of the competition. The mean (-0.637) across all replicates from all densities and inoculum ratios for the center was used as the baseline advantage of ΔwbpL over ΔwapR. This difference in advantage was subtracted from all competitions involving strains with these LPS biosynthesis gene deletions.

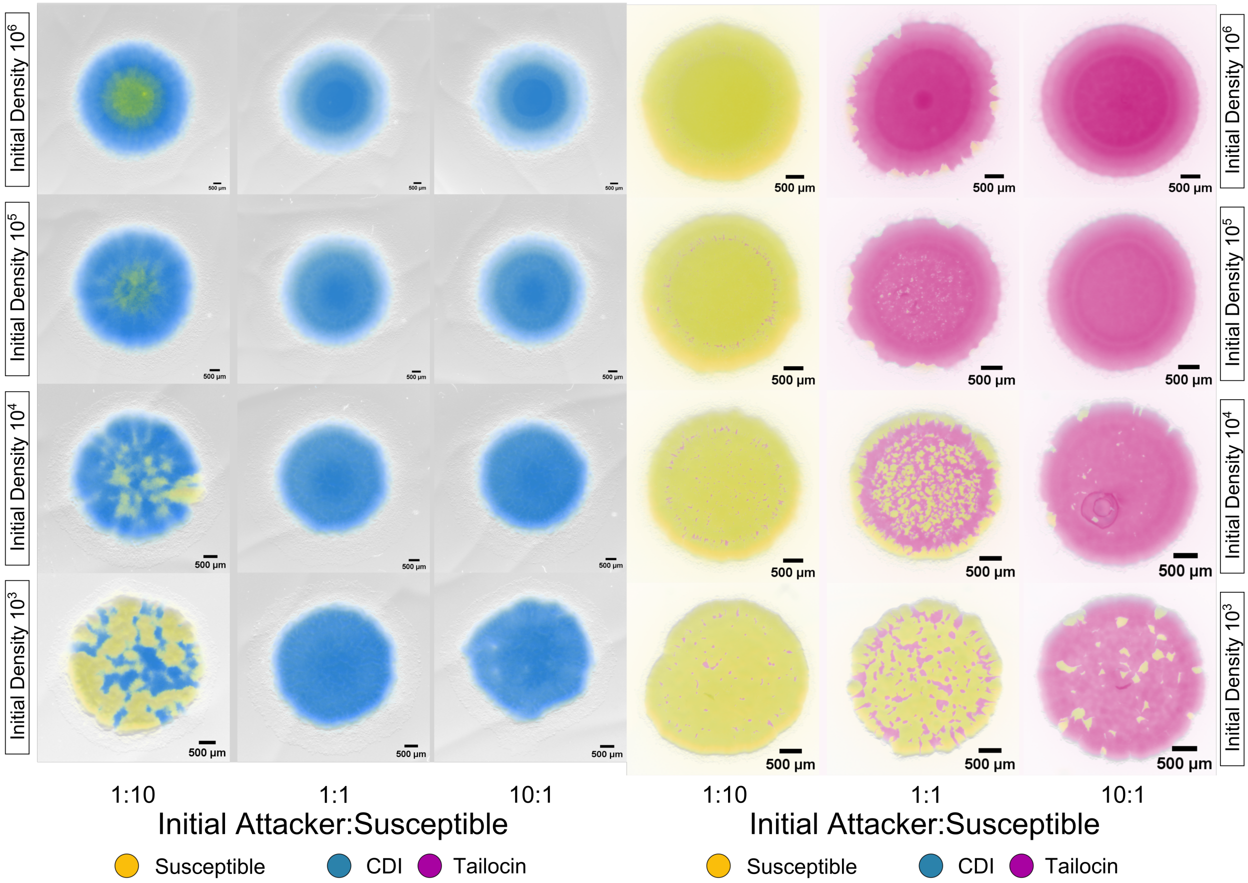


**Figure S5: Colony competitions of contact dependent inhibition (CDI) and tailocins highlight differences between contact and diffusible toxins**. Representative microscopy images of colony competitions inoculated from different starting densities (mean inoculum density 1.9 * $10^3$, $10^4$, $10^5$, $10^6$ CFU/µL). and initial ratios of attacker to susceptible cells taken after 48 h of growth. All strains are expressing constitutive fluorescent protein genes and false-coloured either blue (CDI attacker, top), magenta (tailocin attacker, bottom) or yellow (susceptible, top and bottom). Scale bar indicates 500 µm. For the CDI competitions the attacker was wild-type and the susceptible has the CDI toxin and anti-toxin deleted. For the tailocin competitions, the attacker is ΔwapR and the susceptible strain is ΔR2ΔwbpL.

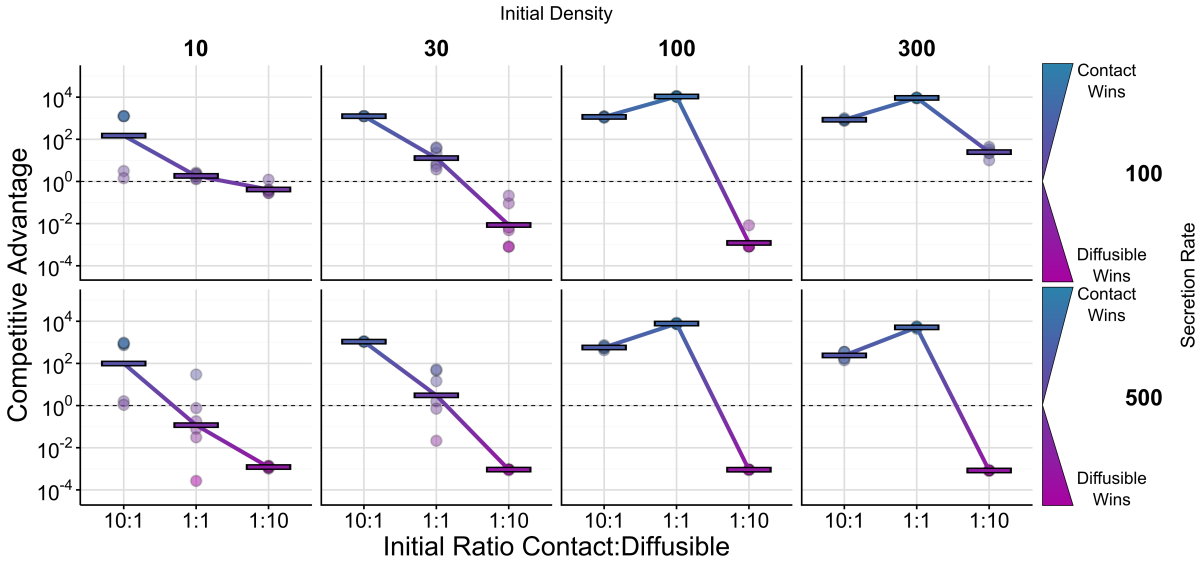


**Figure S6: Agent-based modelling of head-to-head weapon competitions between short and long-range weapon users.** Quantification of simulated direct weapon competition outcomes started at different initial densities, ratios and secretion rates. Density indicates the initial number of cells in the simulation. Competitive advantage assesses the log fold change in the attacker strain compared to its competitor from the beginning to end of the competition. Densities of 10 and 100 cells, with secretion rate 100, correspond respectively to “Low” and “High” starting densities shown in Figure 3.

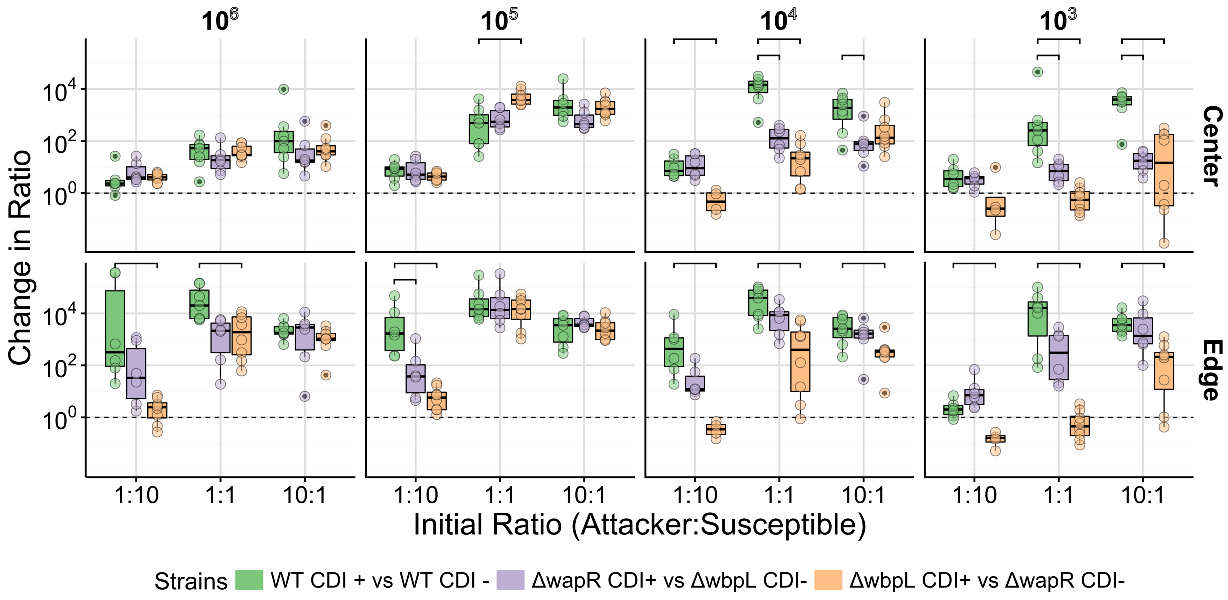


**Figure S7: Outcomes of colony competitions shows that CDI remains functional in LPS biosynthesis gene mutants, but its effectiveness is diminished at low densities.** Outcomes of CDI mediated competitions in wild-type (WT, green) or asymmetric LPS backgrounds. For these cases, both strains also have tailocins (pyocin R2) deleted. The attacking CDI+ strain is either ΔwapR against a CDI susceptible ΔwbpL (mauve) or CDI+ ΔwbpL against CDI susceptible ΔwapR (orange). Colonies were inoculated at the denoted initial densities (mean inoculum density 2.0 * $10^3$, $10^4$, $10^5$, $10^6$ CFU/μL). and quantified by sampling, plating and counting colony forming units after 48h of growth. Competitive advantage assesses the log fold change in the attacker strain compared to its competitor from the beginning to end of the competition. Top brackets indicate a significant difference between each single weapon and the combination of weapons (two-sided Welch's t-test, $p < 0.05$, Benjamini-Hochberg MHT corrected 0.95).

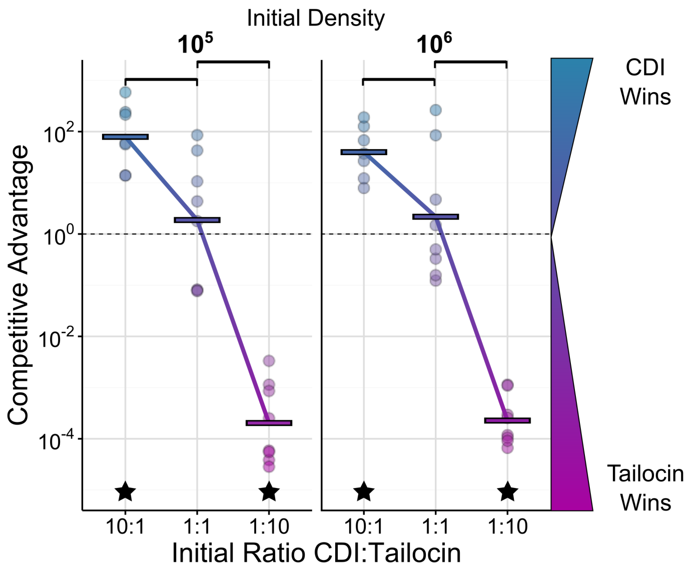


**Figure S8: Head-to-head weapon competitions between short and long-range weapon users at the colony edge.** Quantification of direct weapon colony competition outcomes at the colony edge by sampling, plating and counting colony forming units. Competitive advantage assesses the log fold change in the attacker strain compared to its competitor from the beginning to end of the competition. Values above 1 ($10^0$, dashed line) indicate an advantage for CDI, while values below 1 (i.e. $10^{-2}$, $10^{-4}$) indicate an advantage for tailocins. Lines indicate the mean of replicates ($n \geq 6$). Top brackets indicate a significant difference between the initial ratios (two-sided Welch's t-test, $p < 0.05$, Benjamini-Hochberg MHT corrected 0.95). Stars indicate a significant competitive advantage (one-sided Welch's t-test, $p < 0.05$, Benjamini-Hochberg MHT corrected 0.95). Competitions were inoculated with different initial densities (mean inoculum density 2.3 * $10^5$, $10^6$ CFU/µL). The genotype of the CDI using, tailocin susceptible strain (blue) is ΔR2ΔwbpL. The genotype of the tailocin using, CDI susceptible strain is ΔwapRΔCDI.

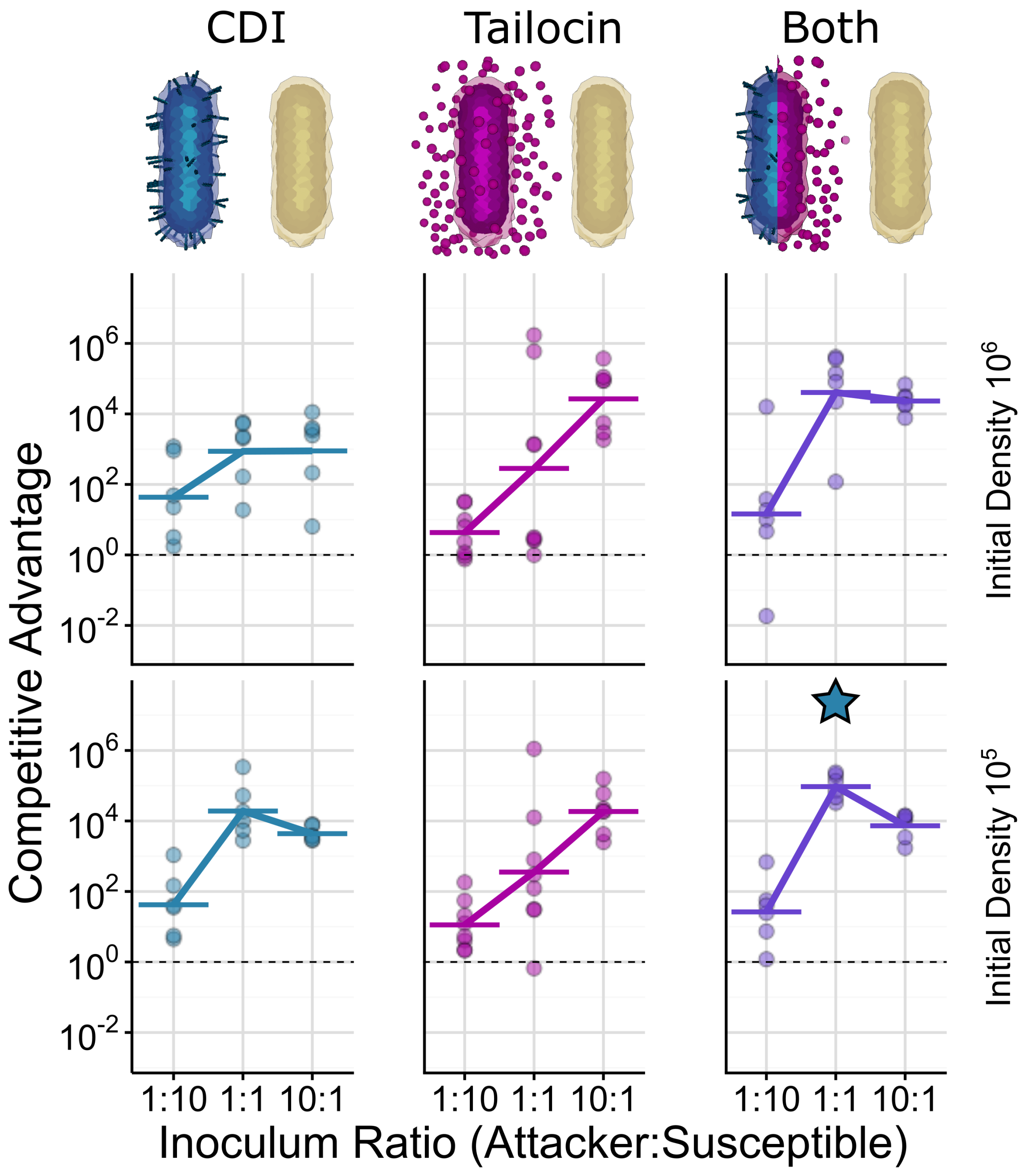


**Figure S9: Short and long-range weapon benefits can combine positively at the colony edge.** Quantification of competition outcomes in the colony edge for two initial cell densities (mean inoculum density 1.9 * $10^5$, $10^6$ CFU/μL). Competitive advantage assesses the log fold change in the attacker strain compared to its competitor from the beginning to end of the competition. Competitions where the attacker has just CDI (blue, left), just tailocins (magenta, centre) or both weapons (purple, right) show the advantage gained from using two weapons together as compared to just one. Data are adjusted to account for differences in competitiveness of the strain backgrounds (ΔwapR relative to ΔwbpL; see methods and Figure S2). Horizontal lines indicate the mean of replicates ($n \geq 6$). The star above the double weapon data indicates a significant difference between the combination of weapons and CDI (two-sided Welch's t-test, $p < 0.05$, Benjamini-Hochberg MHT corrected 0.95). Data from colony edge are noisier than in the colony center but patterns are consistent with the colony interior.

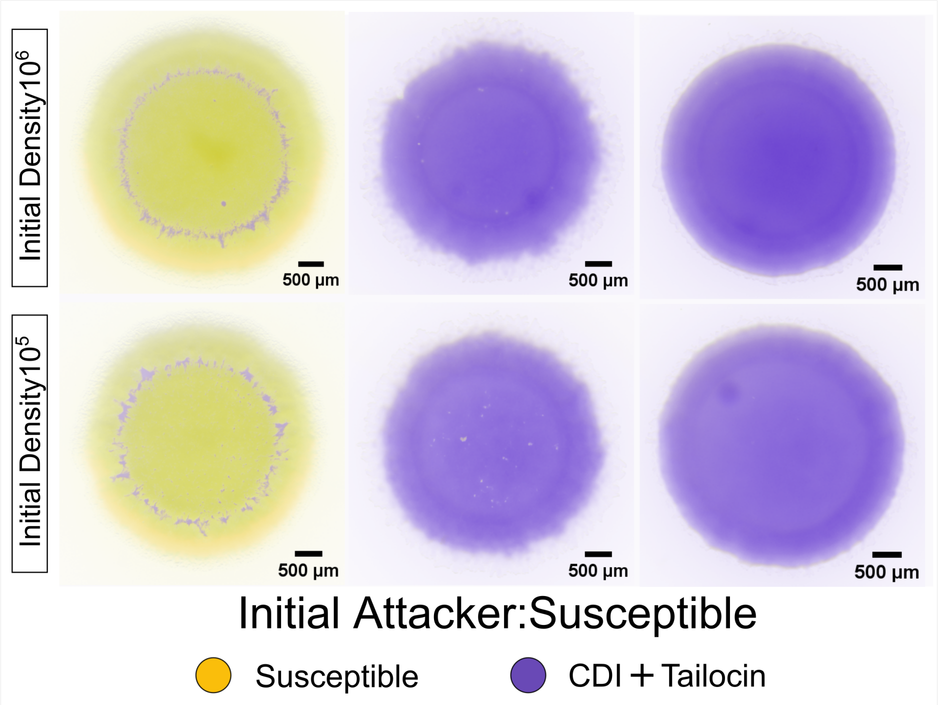


**Figure S10: Microscopy images of colony competitions with doubly-armed attackers.** Representative microscopy images (taken after 48 h of growth) of colony competitions inoculated from different starting densities (mean inoculum density 1.9 * $10^5$, $10^6$ CFU/µL). and initial ratios of attacking and dual CDI/tailocin susceptible cells. All strains are expressing constitutive fluorescent protein genes and false-coloured either purple (attacker) or yellow (CDI and tailocin susceptible). Scale bar indicates 500 µm. The genotype of the attacker is Δ*wapR*. The genotype of the susceptible strain is ΔCDIΔR2Δ*wbpL*.
